# Dynamic Structure Of Motor Cortical Neuron Co-Activity Carries Behaviorally Relevant Information

**DOI:** 10.1101/2022.05.18.492501

**Authors:** Marina Sundiang, Nicholas G. Hatsopoulos, Jason N. MacLean

**Author notes:** co-senior authors (Nicholas G. Hatsopoulos, Jason N. MacLean). Correspondence: Jason N. Maclean, Nicholas G. Hatsopoulos.

## Abstract

Skillful, voluntary movements are underpinned by computations performed by networks of interconnected neurons in the primary motor cortex (M1). Computations are reflected by patterns of co-activity between neurons. Using spike time correlations, co-activity can be represented as *functional networks (FNs)*. Here, we show that the structure of FNs constructed from instructed-delay reach trials in non-human primates are behaviorally specific: low dimensional embedding and graph alignment scores show that FNs constructed from closer target reach distances are also closer in network space. We next constructed *temporal FNs* using short intervals across a trial. We find that temporal FNs traverse a low-dimensional subspace in a reach-specific trajectory. Alignment scores show that FNs become separable and correspondingly, decodable shortly after the instruction cue. Finally, we observe that reciprocal connections in FNs transiently decrease following the instruction cue, suggesting the network momentarily switches from a recurrent system to one that is more feedforward.

## INTRODUCTION

Individual neurons do not function in isolation, but rather as co-active and cooperatively interacting components of spiking networks.The resulting coordinated neuronal co-activity is reflected in pairwise spike time correlations. Spike time correlations have been implicated in the propagation of spiking in networks (Bojanek et al., 2020; Chambers and MacLean, 2016) making them an intriguing signal when studying the circuit mechanisms and computations that give rise to behavior. Recent work has shown that spike time correlations are an informative description of visual stimuli (Dechery and MacLean, 2018; Kotekal and MacLean, 2020; Levy et al., 2020; Ruda et al., 2020), auditory stimuli (Betzel et al., 2019; Insanally et al., 2019), and spatial location (Levy et al., 2022; Nardin et al., 2021). In motor cortex (M1) pairwise spike count correlations have been shown to provide information about motor behavior beyond what is provided by firing rates alone (Maynard et al., 1999), and have also been used to improve encoding models that predict the activity of neurons (Stevenson et al., 2012). However, the temporal dynamics of spike time correlations over the course of a behavioral trial as well as the extent of specificity of spike time correlations to kinematic variables is underexplored.

Here, we represent pairwise spike time correlations as *Functional Networks (FN)* and use tools from network science (Bassett and Sporns, 2017; Bullmore and Sporns, 2009; Rubinov and Sporns, 2010) to examine the structure and information content of spike time correlated activity of active populations in M1. Given that many of the computations needed to carry out a movement happen in fractions of a second, averaging interactions over long periods of time are unlikely to capture important aspects of network structure, such as how and when the network architecture changes in order to support these computations (Pedreschi et al., 2020). We extend the FN framework by constructing temporal FNs (Holme and Saramäki, 2012; Ju and Bassett, 2020; Thompson et al., 2017) of single trials by computing spike time pairwise correlations across short intervals to elucidate the temporal progression of pairwise correlations among a population of M1 neurons throughout a trial and to relate single-trial population dynamics to network structure.

Using temporal FNs, we evaluate whether the dynamics of pairwise correlations carry information about the instructed reach, and when network structure begins to carry that information relative to behaviorally relevant moments during the trial. Moreover, we determine what types of network interactions are most prevalent relative to different phases of the trial. We find that temporal FNs evolve during the trial in a way that specifies the instructed reach. Using a perceptron decoder, we demonstrate that temporal FNs carry information about motor behavior beyond what is conveyed by firing rate changes alone. Finally, we show that the topology of temporal FNs varies systematically over the time course of a trial, becoming transiently less reciprocal after the onset of the instruction cue in an instructed-delay movement task, suggesting that FNs switch from a highly recurrent system to one that is more feedforward at this key moment in the trial.

## RESULTS

We analyzed single-unit recordings from M1 while two macaques (Monkey Rs and Monkey Rj) performed a center-out reaching task in the horizontal plane. Subjects were trained to control a cursor on a screen by moving a handle on a 2D arm exoskeleton (KINARM, Kingston, Ontario). The task was to hold the cursor in the center position during an instructed delay period while an instruction cue was presented that indicated one of 8 target reach locations positioned radially around the center. After the end of the delay period (1s), a go cue appeared instructing the subjects to initiate a reaching movement (Figure 1A).

**Figure 1.**
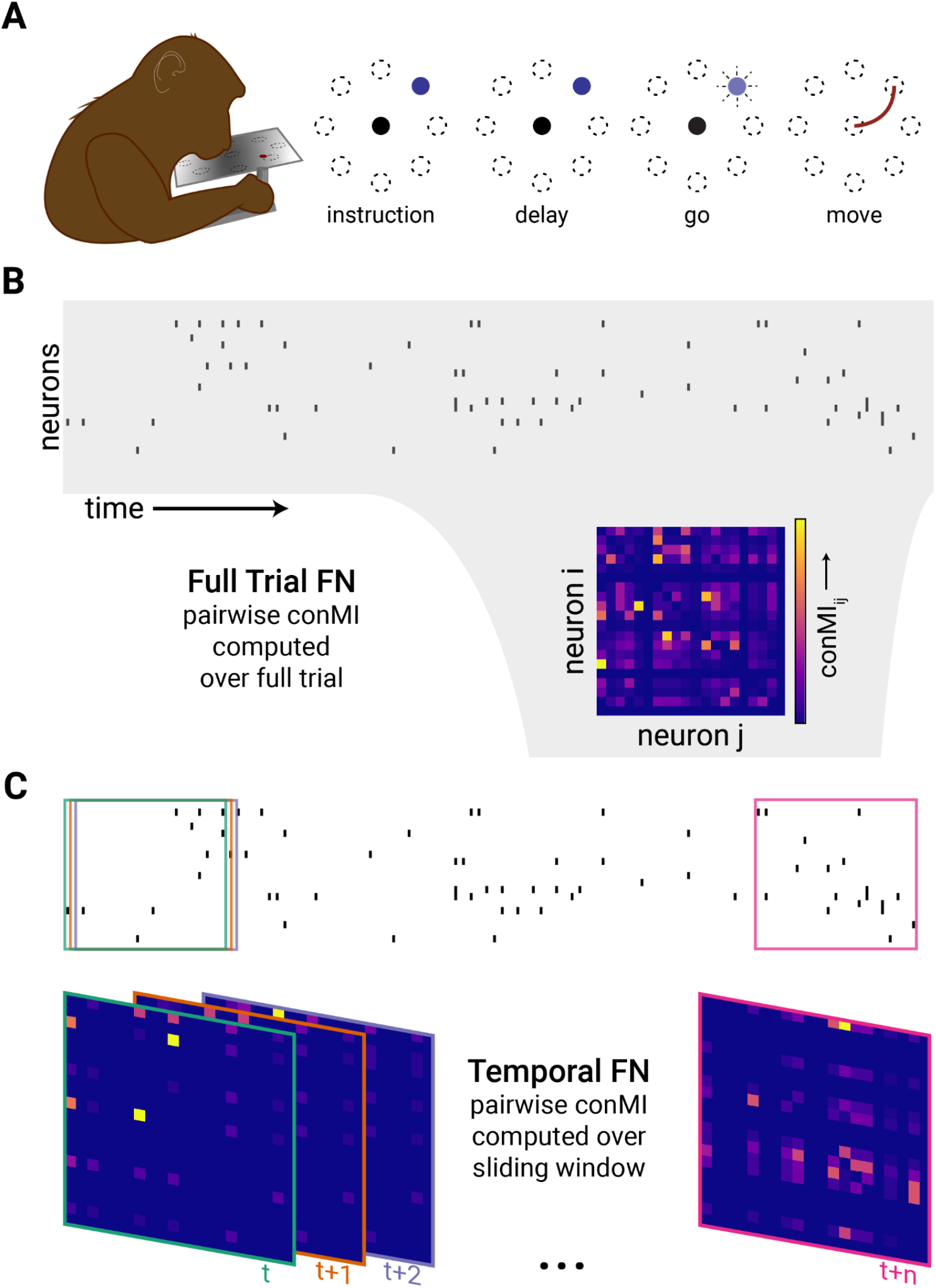
Task structure and Functional Network construction. **A)** Subject using a Kinarm exoskeleton to control a cursor on a screen. The subject holds the cursor at the center target (black circle) to initiate a trial. An instruction cue (blue) appears in one of 8 target directions (locations of targets are indicated by the circles with broken-lines). After the delay period, the target cue begins blinking which indicates that the subject may move to the instructed target. **B)** A raster plot shows the binned spikes of the recorded neurons across time. The Full Trial FN is constructed by computing the confluent Mutual Information (conMI, see Methods) between the spike train for the full trial for each pair of recorded neurons. **C)** To generate the Temporal FN, pairwise spike correlations were computed within a 200 ms sliding window across the trial. Boxes on the raster indicate the interval in the spike train that was considered for the corresponding FN in the with the same color border.

We summarized neuronal population activity during the task as a functional network (FN, Figure 1B-C). The FN is composed of nodes, which are the spiking single units, and edges between nodes, which correspond to the pairwise spike time correlations and not anatomical connectivity (Rs: 142 single units, Rj: 78 single units). Specifically, edges are computed using a mutual information measure (conMI, Methods; Chambers et al., 2018). Confluent mutual information (conMI) computes the mutual information between the binned activity of neuron i at time t, and of neuron j at time t and the adjacent time bin, t+1 (where bins are 10ms) and is positive, by definition. The reliability of the pairwise correlation between neuron i and j during a specified time window resulted in a weight. The resultant FN is both weighted and directed corresponding to the reliability and the directionality of the pairwise spike time correlation between units.

### Correlational structure in population activity are specific to reach direction

We first determined whether FNs computed from pairwise spike time correlations during the entirety of single reach trials are specific to reach target direction. We did so in two ways: low-dimensional embedding of FNs and a graph similarity metric. Both approaches show that FNs constructed from closer reach target distances are also closer in network space. First, we embedded the FNs in a low dimensional manifold that optimizes the distances between data points according to the original high-dimensional similarity. Consequently, FNs that are more similar to each other will lie closer together on the low dimensional manifold, and conversely, FNs that are structurally different will be more distant. Specifically, we used UMAP to project the high dimensional FNs into a lower dimensional space (see Methods). We found that networks from reaches to neighboring directions are close together in the low-dimensional manifold. In fact, the projection produced a radial arrangement of data points that reflected the radial arrangement of the target locations (Figure 2A-D). We note that monkey Rj’s kinematic trajectories towards the lower right targets were not straight toward the target (pink in Figure 2C); Rj moves straight down and then right. We found that FN embeddings where the target is the lower right (pink) and the one straight down (violet), are correspondingly overlapping.

**Figure 2.**
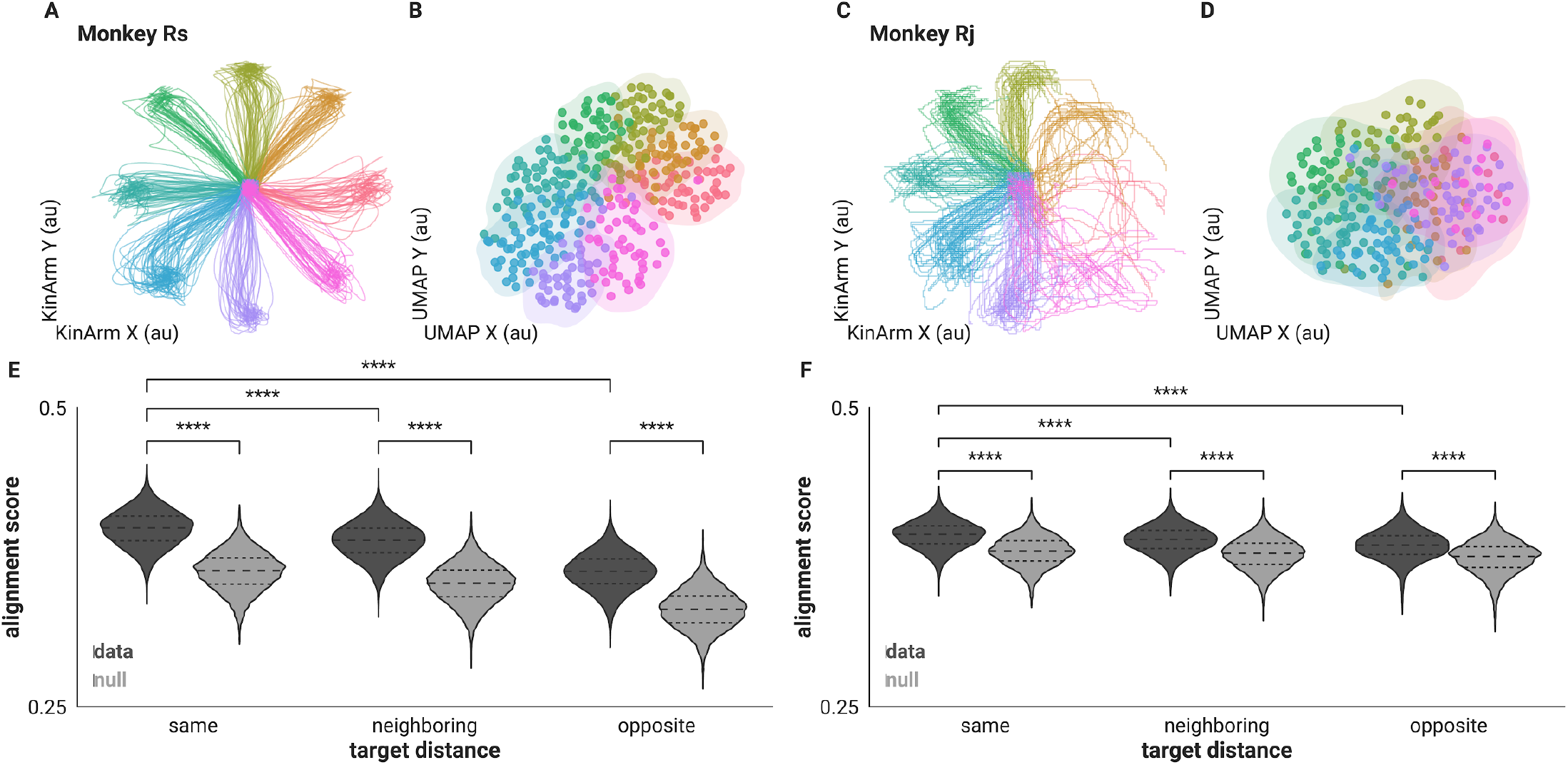
Correlational structure of full trial FNs are specific to reach target direction. **A**,**C)** Hand trajectories of included trials colored by reach direction. **B**,**D)** Low-dimensional projection of full trial FNs preserve reach target direction. Colored the same as A,C. Shaded contour is the bivariate Kernel Density Estimation (see Methods) for the (x,y) positions of the FNs to visualize the boundaries of the projected FNs of each direction. **E**,**F)** Distribution of alignment scores between each pair of FNs separated by target distance in dark gray and score distributions for corresponding rate-matched FNs shown in light gray (**** = p < 1.00e-04, Mann-Whitney U two-sided test). Dashed lines indicate quartiles.

To further evaluate the similarities and differences between FNs according to reach direction we compared networks using Graph Alignment Scores (GAS, see Methods). GAS allows for a quantitative comparison of FNs by identifying common edges between graphs (Gemmetto et al., 2016). This metric preserves node identities and reflects the fraction of similar edges out of all the existing edges between two networks. Therefore, a value of 1 means that the two networks are exactly the same. We measured alignment between each pair of FNs constructed from trials of different reach directions. We then grouped the measured GAS according to the degree difference between the respective instructed reach targets: same (Δ0°), neighboring (Δ45°) and opposite (Δ180°). We observe that pairs of FNs from trials with the same reach target have higher alignment scores (Δ0° Rs = 0.399 ± 0.014, Rj = 0.393 ± 0.011, mean ± std) than those from neighboring reach directions (Δ45: Rs = 0.388 ± 0.015, Rj = 0.389 ± 0.011; p<=1.865e-89, Mann-Whitney U (MWU) two-sided test, mean ± std). Alignment scores were lowest when computed using FNs from opposite reach targets (Δ180°: Rs = 0.363 ± 0.015, Rj = 0.385 ± 0.012; p<=1.882e-236, MWU two-sided test, mean ± std) indicating that the shared correlational network structure, in this case summarized as FNs, are informative of the instructed reach (Figure 2E-F). Additionally, compared to the rate matched networks (see Methods), GAS from data were higher for all three distributions suggesting that trial-to-trial pairwise spike time correlations in the data are not replicated by the rate-matched networks (Rs = 0.363 ± 0.016, 0.353 ± 0.016, 0.331 ± 0.016 ; Rj = 0.380 ± 0.012, 0.378 ± 0.012, 0.374 ± 0.014; GAS for pairs of rate-matched FNs for same, neighboring, and opposite trials, respectively, p<0.001, MWU two-sided test for all data and rate-matched comparisons, mean ± std).

Together both measures, UMAP and GAS, demonstrated that the pairwise spike time correlation structure within the recorded population is specific for each reach direction, and that these differences were not simply the consequence of the firing rate differences.

### Temporal progression of pairwise spike time correlations on single trials are specific to reach direction

Correlations between neurons are dynamic (Aertsen et al., 1989) and can vary over a behavioral trial depending on task condition (Vaadia et al., 1995). Thus, we next evaluated the time varying spike time correlational structure by constructing FNs from short epochs (200 ms) of trial time. These “snapshots” of the network across time, *temporal FNs*, allowed us to evaluate how the network evolves over the time course of single trials and to determine when, during a trial, the reach-specific differences in the FNs first occur and are most pronounced. Again we examined the differences between FNs in two ways. First, we projected the FNs into a lower-dimensional subspace using UMAP as we had done with the full trial FNs. This allowed us to use the projection to track the progression of temporal FNs relative to trial structure and reach direction. We found that the temporal FNs progressed along trajectories through the subspace in a reach-specific path (Figure 3); Temporal FNs are initially close together in the computed subspace (Figure 3A) and then separate according to the instructed reach direction as the trial progresses (Figure 3B), separating maximally after the *go* cue and during movement (Figure 3C). Again, we used an unsupervised method of dimensionality reduction and therefore did not include information about the behavior of the subject in the projection. Consequently, any separation in the projection of the temporal FNs is the result of the differences in the pairwise spike time correlations of the neural population and not from any training labels.

**Figure 3.**
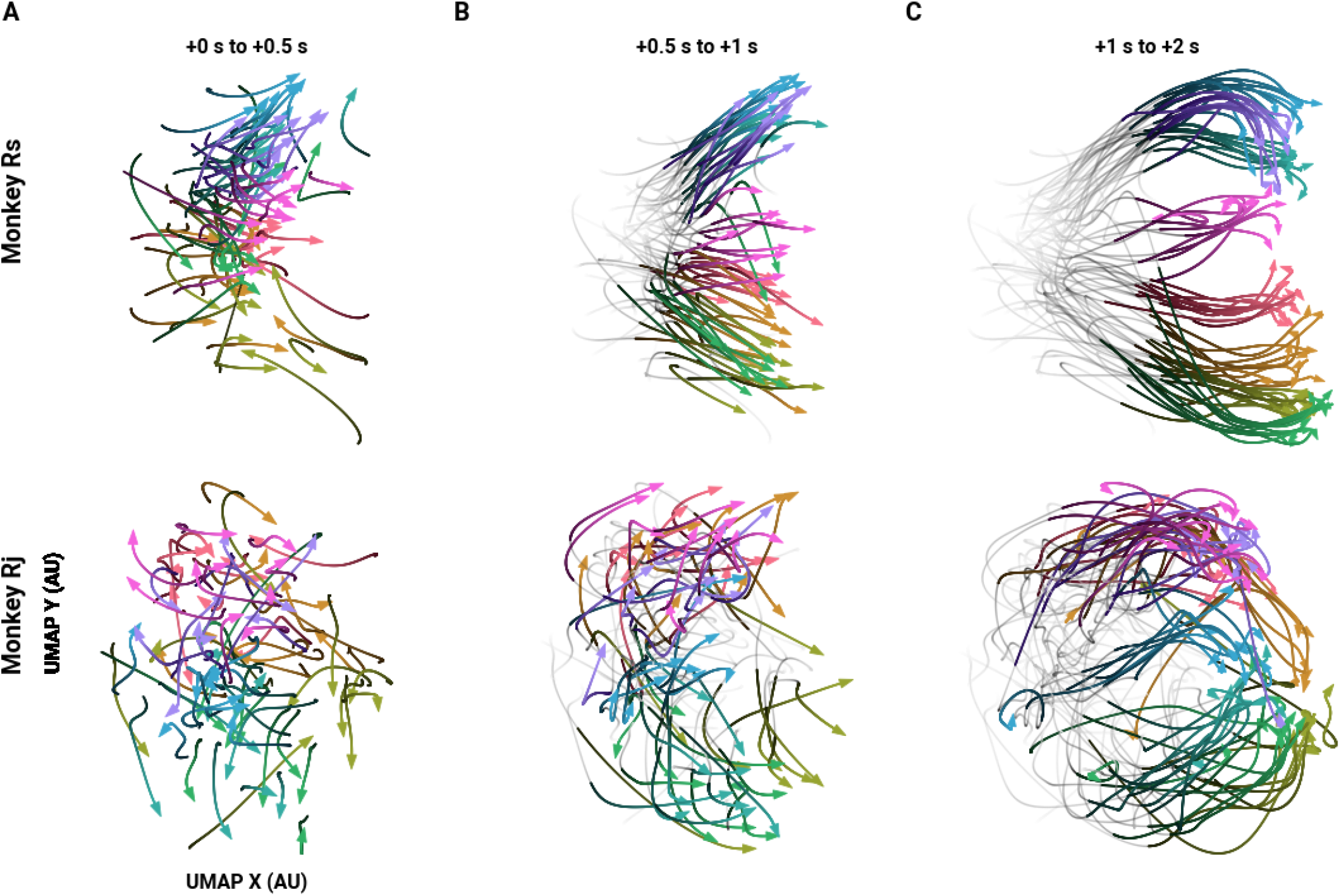
Trajectories of single trial temporal FNs through a low-dimensional subspace. Example trajectories of single trial temporal FNs across a low-dimensional subspace (10 trials per direction). Times (in seconds) indicate the time post-instruction. **A)** Trajectories from instruction (0s) to 0.5 s after instruction. Trajectory begins with the darkest hue. Color legend the same as Figure 2a-d. **B)** The same trajectories from (A) later in time: the gray, low-opacity tails show the trajectory from instruction, and the colored and opaque section show the trajectory at 0.5 s post-instruction to Go cue (1s post-instruction). **C)** the same as (B) but for after Go (1 s) and through the end of the trial.

As with the full trial FNs, we next computed the extent to which the structure of the temporal FNs indicated the instructed reach. At each time window, we measured the GAS of every pair of FNs and sorted the scores according to difference in target direction. We then evaluated the distribution of graph alignment scores for the same, neighboring, and opposite reach directions across the time course of the trial (Figure 4). Before and shortly after instruction, the FNs are not informative as to reach direction as indicated by similar GAS values (MWU two sided test, p >= 4.045e-3, Bonferroni corrected between all scores from data FNs). The score distributions of neighboring and opposite reach-directions became significantly different from scores of FNs from the same reach trials post-instruction cue consistent with the embedding result (Figure 4A; Δ0° vs Δ180°: MWU two sided test, p <= 1.576e-6, Bonferroni corrected from 170 ms post instruction for Rs, and 180 ms post instruction for Rj. Δ0° vs Δ45°: MWU two sided test, p <= 3.106e-4, Bonferroni corrected from 210 ms post instruction for Rs, and 270 s post instruction for Rj). The scores remained significantly different for the remainder of the trial and also at movement onset (Figure 4B-C). Notably, the scores between the same and the opposite reach directions diverge first before scores from the same and neighboring reach directions, again suggesting that FNs generated from nearby reach directions are also more similar in network space.

**Figure 4.**
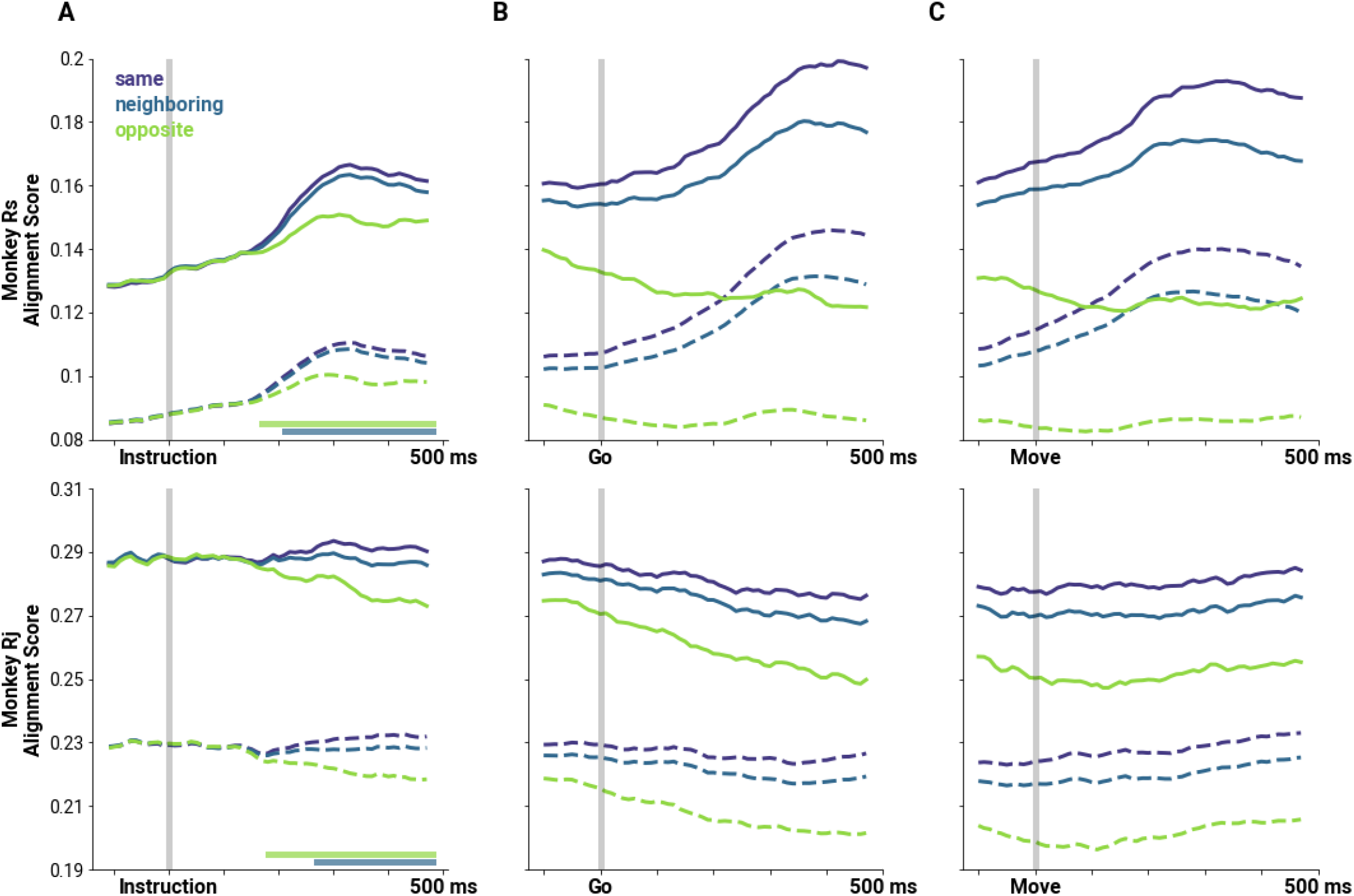
Temporal progression of correlational structure on single trials are specific to reach target direction. **A)** Mean alignment scores between pairs of FNs at each sliding window (solid lines), and corresponding scores for rate-matched FNs (broken lines) aligned to the instruction cue. Alignment scores are separated based on reach target difference: same (violet), neighboring (blue), and opposite (green) reach targets. Straight lines on the lower left of the plot indicate when the score distributions from neighboring or opposite directions (blue and green, respectively) become significantly different from the score distributions between FNs of the same direction (p<0.001 MWU two-sided test, Bonferroni corrected). **B)** same as (A) but aligned to Go cue. Differences between distributions are significant throughout the plotted window (p<0.001 MWU two-sided test, Bonferroni corrected). **C)** same as (A) but aligned to movement onset. Differences between distributions are significant throughout the plotted window (p<0.001 MWU two-sided test, Bonferroni corrected).

### Decoding behavior from functional networks

Temporal FNs became increasingly differentiable during the trial with the differences following a stereotyped time course relative to trial structure, indicating that there is information about motor behavior in the temporal FN. We next asked if this information can be used to decode reach target direction. Since GAS analysis indicated that the differences in the temporal FN depended on the time of comparison, we also measured when the temporal FNs were decodable relative to trial structure and movement onset, and when they were maximally decodable. To do so, we employed multilayer perceptron decoders trained on different features of the data to predict the reach target of a 200 ms sample window of a single trial (see Methods). We then compared the performance of the decoders based on the features on which they were trained. Specifically, decoders were trained on either the set of pairwise spike time correlations between neurons generated in the time window (FN decoder, Figure 5 green), or the firing rates of the neurons during the window (FR decoder, Figure 5, blue). To determine whether the correlations carry non-redundant information beyond firing rates, we also examined decoding performance when both firing rate and the temporal FNs were included in decoder training (FRFN decoder, Figure 5, red). Comparing the decoding performance of the FN and FR decoders to the full FRFN decoder allowed us to establish the contribution of each held out feature to the overall decoding performance. Finally, in order to isolate the importance of the data specific correlations rather than correlations that arose from the varying firing rates of the recorded neurons, we generated FNs from Poisson rate matched neurons and similarly evaluated decoding performance (null FN decoder, Supplementary Figure 1, violet; see Methods).

**Figure 5.**
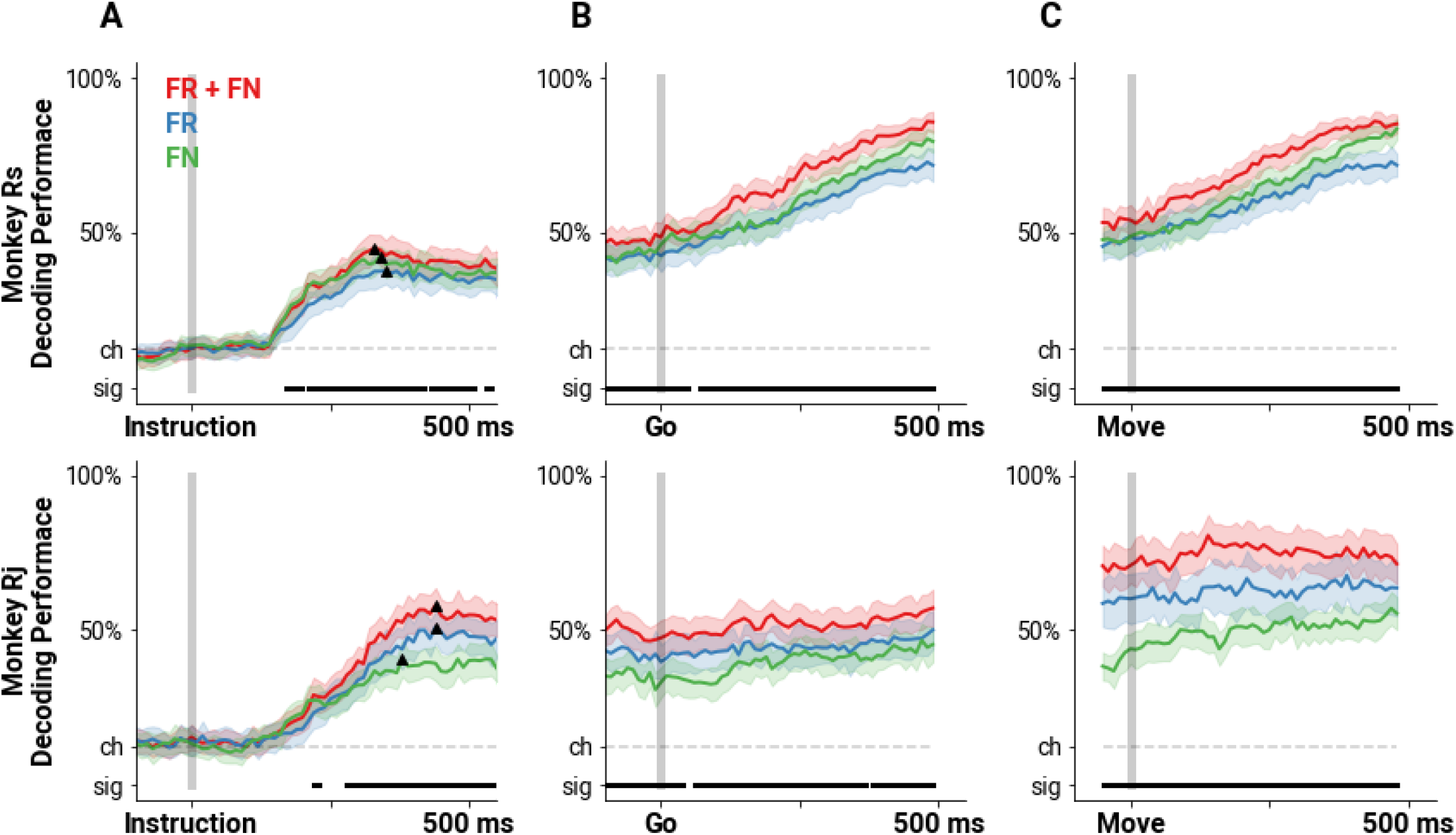
Decoders that incorporate pairwise spike time correlations predict reach direction more accurately. Mean performance perceptron decoders trained on either firing rates (FR, blue), the set of pairwise correlations as FNs (FN, green) or both firing rates and correlations (FR + FN, red) aligned on Instruction cue (**A**), Go cue (**B**), and movement onset (**C**). Dashed gray line indicates chance level (1 in 8), and the bar on the bottom of each panel indicates the times during which performance of the FR and FRFN decoders are significantly different from each other (p<0.001 MWU two-sided test, Bonferroni corrected). The triangles in panel A mark the initial peak in performance for the decoders. The shading represents standard deviation of the decoder performance.

As expected, before the instruction cue, all four decoders performed at chance level (Figure 5 and Supplementary Figure 1, 12.5% performance indicated by dashed line). At 147 ± 9 ms following the instruction cue, decoding performance increased above chance for the FN, FR and FRFN decoders (1 sample t-test, p <= 9.550e-4, Bonferroni corrected, mean ± std). The null FN decoders achieved performance above chance about 100 ms later, at 240 ± 10 ms (mean ± std). All four decoders reached an initial peak around 364 ± 48 ms (defined as the maximum performance within 500 ms of the instruction, indicated by the black triangle on every performance trace in Figures 5 and 6). At this peak, the FRFN decoders performed better than the FR decoder (Rs: 45.10% vs 37.67%, Rj: 58.03% vs 51.03%, respectively, p <= 6.438e-05, MWU two-sided test), suggesting that the inclusion of pairwise spike time correlations provides additional information about the reach direction. In fact, the FRFN decoder performance was significantly higher than the performance of the FR decoder beginning 170 ms for Rs and 220 ms for Rj and remained significantly higher throughout the rest of the trial, as well as during movement (MWU two sided test, p <= 9.888e-04, Bonferroni corrected between FRFN and FR performance scores). We also observed that at the initial peak, the FN decoders performed significantly better than the rate-matched FNs (Rs: 42.39% vs 33.37%, Rj: 40.85% vs 27.28%, respectively, p<=3.562e-04, MWU two-sided test). For Rj, this performance difference was significant from 170 ms throughout the rest of the trial (MWU two sided test, p<=3.3845e-04, Bonferroni corrected). For Rs this difference was only significant early in the delay period (310 to 390 ms, MWU two sided test, p<=8.853e-04, Bonferroni corrected) and sporadically after the go cue. This highlights that there is information about the instructed reach in the pairwise spike time correlations in the data that does not arise from chance correlations due to firing rates.

**Figure 6.**
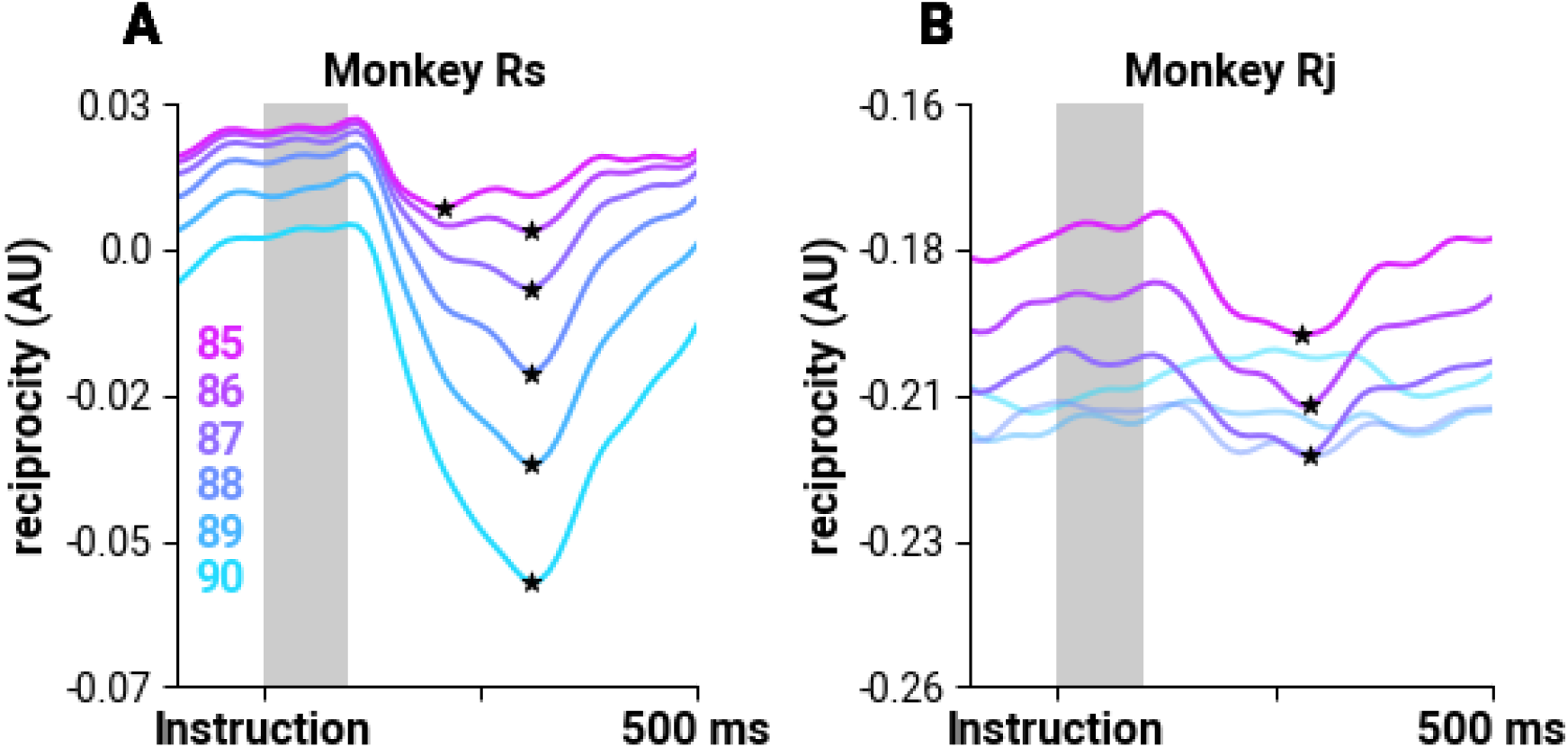
Reciprocity decreases shortly after instruction. Mean normalized reciprocity scores from different threshold values for monkeys Rs (**A**) and Rj (**B**). The mean reciprocity for the shaded region (Instruction to 100 ms post instruction) is computed for each trial resulting in a distribution of reciprocity scores, which is then compared to the scores at the minima to determine if there is a decrease in reciprocity. Stars and full opacity lines indicate whether the distributions at the minima decrease significantly (p<0.001, independent sample T-test)

### Reciprocity in the network varies systematically during the trial

One of the strengths of representing cortical population activity as a weighted and directed temporal FN, is that not only can we investigate the dynamics of pairwise spike time correlations across the population (as edge weights), we can also consider the changes to network architecture (as edge direction). Doing so has the potential to provide insights into information propagation and computation. Here, we focus on the most reliable reciprocal connections in the temporal FN. We thresholded the temporal FNs, according to weight, in order to isolate these edges and then measured the normalized reciprocity of the temporal FNs throughout the trial (see Methods). Reciprocity tells us how likely two units in the temporal FN are mutually linked (Squartini et al., 2013), and serves as a building block for higher-order structures such as triplet motifs (Recanatesi et al., 2019). Additionally, reciprocity also tells us the importance of the direction of the interaction between neurons: a fully reciprocal FN means that units are influencing each other symmetrically in contrast to a less reciprocal FN where the direction of the interaction is important in characterizing the network. In the temporal regime, reciprocity may influence the state of the FNs in the next time point, whether to sustain the interactions or to reorganize. We investigated whether the reciprocity of the temporal FN changes when external inputs arrive at M1. We compared the distribution of reciprocity scores at the time when the mean reciprocity over trials is at its minimum (Figure 6, indicated by ★) within the instructed delay period with the distribution of mean scores of each trial from instruction to 100 ms post-instruction (Figure 6, gray shaded region). We found that reciprocity significantly decreased for the networks for both Rs and Rj (Figure 6, independent sample t-test, p <= 3.773e-4). The minima occurred at 291 ± 31 ms after instruction cue (mean ± std). This suggests that at moments when inputs external to the recorded population would be most likely, the network structure becomes less recurrent and more feed-forward.

## DISCUSSION

We show that pairwise spike time correlations between neurons, summarized as Functional Networks (FNs), are informative of reach direction and are dynamic over the course of a single task trial. Previous studies have shown that interactions between neurons provide additional information about behavior or the environment that is not gleaned from merely observing each neuron independently (Maynard et al., 1999). Moreover, the dynamics of pairwise correlations are indicative of behavioral conditions that are otherwise unobservable from individual neuron firing rates or from trial-averaged cross-correlations (Vaadia et al., 1995). These previous studies measured correlations of spike counts across trials and over broad time scales (see, however, Hatsopoulos et al., 1998 that examined precise directional information in synchrony). We build on this work by examining pairwise spike time correlations and find that FNs computed from a full trial are specific to motor behavior. Since movement is the culmination of multiple computations, many of which happen at short timescales, we hypothesized that information about environmental and sensory cues relevant to the task must be represented in the short time scale pairwise spike time correlations. Our work substantiates this hypothesis, since we find evidence that temporal FNs constructed from short intervals during single trials carry a representation of the instructed movement. Specifically, we found that the similarity between pairs of temporal FNs from single trials reflected the distance between their respective target directions and are correspondingly decodable. The temporal FN framework also allows us to identify *when* correlations begin to be indicative of reach, which in turn has the potential to provide insight into corresponding computations. We find that FNs become decodable above chance 140 ± 10 ms after the instruction cue appeared. Notably an increase in timing precision of spiking in individual neurons across trials has been found around this time (∼100 ms), and mutual information increases between target location and spike counts of single neurons at this time (Reimer and Hatsopoulos, 2010). The modulation of firing rates across the population and the stronger relationship between spike count and target location suggests that information about the target arrives at M1 at this time lag. Consistently, we find that precise spike timing *between* neurons in single trials, as reflected in their correlation, is also informative of target direction. Moreover, alignment scores between temporal FNs constructed from single trials of the same target direction are higher than alignment between temporal FNs that correspond to neighboring and opposite directions, suggesting that there is consistent and informative spike timing between neurons on a trial-to-trial basis that is specific to reach direction.

When external inputs impinge on local circuitry, they must be integrated into ongoing activity and then propagated and processed across the network. Theoretical work suggests that to propagate spiking a network must systematically cycle between higher order motifs that are dependent on reciprocal connectivity (Bojanek et al., 2020). Here we find that reciprocal connections decreased transiently after the onset of the instruction cue. Thus the networks are transiently more feed-forward (as opposed to feed-back or recurrent) which may be an indication of the network integrating and propagating information about external inputs such as visual stimulus related to the instructed target direction. Reciprocity reaches a minimum just prior to the initial peak in decoding performance suggesting that information external to the recorded population impinges on the network at this time to then be incorporated into ongoing activity. Interestingly, this topology is transient; the network returns to its baseline level of reciprocity, but it is important to note that the information about the reach target is maintained in the FN after this transient topological shift since decodability of the FNs remains stable for the remainder of the delay period. In this work, we focus on reciprocal connections to quantify recurrence, however recurrence in the networks can also manifest in higher-order forms such as in cycle motifs (Chambers and MacLean, 2016). Because higher-order motifs have more steps in recurrence, this may have different implications on information propagation during reaching that is worth exploring in the future. Previous studies have shown that the population activity during the preparatory and the movement phase are dynamic (Churchland et al., 2012; Shenoy et al., 2013; Vyas et al., 2020), that they occupy orthogonal subspaces (Kaufman et al., 2014), and that these subspaces can be linked (Elsayed et al., 2016). Consistent with this work on computation through population dynamics, we show that the patterns of coordinated activity of the neural population evolves during the reach trial in a systematic way. Our findings provide evidence that the network reorganizes to a more feed-forward topology which may facilitate processing external inputs, and setting the state of the population according to the instructed movement and may enable the reported transition from the preparatory subspace to the movement subspace.

In this study we did not try to classify the single units functionally or into putative cell classes. Future work would clearly benefit from data in which individual classes of neurons are considered separately. The FN framework can be extended to include labels on nodes and edges such as cell type, or functional properties (Faskowitz et al., 2022). Notably an impressive census of cell types in motor cortex has recently been published by the The BRAIN Initiative Cell Census Network (BRAIN Initiative Cell Census Network, 2021) highlighting the potential for an analytical approach, such as FNs, that can both incorporate cell information and network dynamics. For example in the visual cortex, the topology of single trial FNs depends on the visual stimulus and untuned neurons act as hubs in the network serving an integral role in stimulus coding (Levy et al., 2020). Previous work has shown that modeling functional connections can predict neural responses in M1 more accurately than canonical tuning curves suggesting that the tuning properties of individual neurons is in part a result of network interactions (Stevenson et al., 2012). Additionally, tuning properties are not stationary such that single cortical neurons in M1 encode temporally extensive movement fragments or trajectories, as opposed to single movement parameters such as direction (Hatsopoulos et al., 2007). The temporal FN framework that we introduce in this paper has the potential to link dynamically changing network interactions across the trial to time-varying preferred movement trajectory tuning.

Aside from carrying information about the instructed reach, we found that FNs appear to carry kinematic information additional to target reach direction. For example, in the full trial and temporal embeddings of FNs from Rj (Figure 2C-D, Figure 3, bottom row), overlapping projected FNs correspond to trials that had similar kinematic trajectories despite different target directions. Additionally, low-dimensional projections of temporal FNs separate into two main branches. Specifically, in Rs, the upper branch corresponds to pulls towards the body. Conversely the lower branch corresponds to reaches that required the subject to push away from the body. This suggests that, while we have performed these analyses on data where the subject is performing an 8-direction center out task, FNs may be able to capture a wider range of movement directions and kinematic variability than explored here.

FNs are a useful tool to achieve insight into M1 network dynamics that correspond to motor behavior. FNs provides a tractable summary of population activity that is decodable and can provide insight into how the state of the system changes over the time course of a trial. The neural interactions summarized by FNs influence spiking activity of the population that can be, in turn, read out by downstream agents (Levy et al., 2020). Our results suggest that FNs reorganize according to motor goals and argue that a functional network examination of motor cortical FNs may provide insights into the underlying circuit mechanisms that drive these cortical population responses and ultimately motor behavior.

## Acknowledgements

We thank Jacob Reimer, Zach Haga, and Dawn Paulsen for collecting the data presented in this work. This work is funded by NIH R01NS111982 and R01NS104898. We thank Mayaan Levy, Caleb Sponheim, Wei Liang, and Gabriella Wheeler Fox for their helpful comments on the manuscript.

## Author Contributions

Conceptualization, M.S., N.G.H, and J.N.M; Methodology, M.S., N.G.H, and J.N.M; Software, M.S.; Investigation, M.S.; Writing - Original Draft, M.S., N.G.H, and J.N.M ; Writing - Review & Editing, M.S., N.G.H, and J.N.M; Visualization, M.S.; Supervision, N.G.H, and J.N.M; Funding acquisition, N.G.H, and J.N.M.

## METHODS

### Lead Contact

Further information and requests for resources should be directed to and will be fulfilled by the lead contact, Marina Sundiang

### Materials availability

This study did not generate new unique reagents.

### Data and Code Availability

Data and custom code to reproduce figures can be made available upon request to the Lead Contact.

### Data Collection and Reaching Task

We used previously published datasets from two macaques performing an instructed center-out reaching task (Hatsopoulos et al., 2004; O’Leary and Hatsopoulos, 2006). Subjects were trained to hold a cursor on a center target presented on a video screen using a 2D arm exoskeleton (KINARM, Kingston, Ontario). One of eight radially positioned peripheral targets was then presented and served as an Instruction cue during which time the subjects were required to keep holding the cursor on the center target. After a 1 second delay period, the peripheral target began blinking (Go cue) instructing the subjects to move the cursor to the peripheral target. (Figure 1A). Trial start was 0.5 s before the instruction cue appeared, and trial termination was 0.5 s after the peripheral target was acquired. Trial inclusion depended upon target acquisition within 1.5s following movement onset. Movement onset is defined as the time when the hand velocity reached 5% of the peak velocity of the movement after the Go cue.

Neural data were recorded from 96-channel Utah arrays implanted in the arm/hand area of primary motor cortex (M1) on the precentral gyrus. Spike waveform snippets sampled at 30 kHz were extracted using a user-defined threshold (Cerebus BlackRock Microsystems) and were sorted into individual units using Offline Sorter (Plexon).

### Using pairwise spike time correlations to construct functional networks

To compute pairwise spike time correlations between recorded neurons, we binned the recorded spike trains into 10 ms bins: assigning a value of 1 if at least 1 spike occurred in that bin, and 0 otherwise. We then used the confluent mutual information (conMI) between the binned spike train (Chambers et al., 2018). conMI tells us how much information we gain from the firing state of a source neuron i at time t about the firing state of target neuron j in the same time bin t and the consecutive bin, t+1:

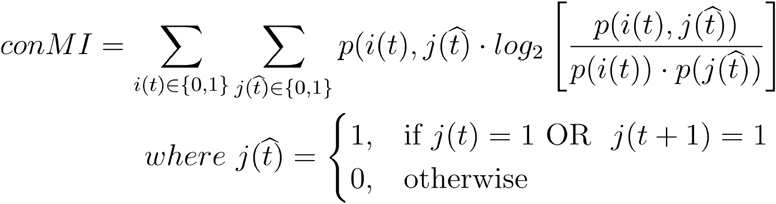

The functional network constructed using this measure is consequently weighted and directed. We computed functional networks from either the full trial duration or from 200ms epochs. For the full trial networks, we computed the conMI between the entire spike train of the source and target neurons for a single trial (Figure 1B). We also computed the conMI for 200 ms windows that we slide in 10ms (1 bin) increments across the full trial duration resulting in a temporal FN: a set of FNs each representing the measured co-activity of the recorded population at each time window (Figure 1C).

To establish if FNs were not simply the consequence of firing rate, we computed FNs from rate-matched Poisson neurons. The rate-matched nulls indicate what structure arises from chance correlations that are due to the firing rates. We computed the instantaneous firing rate of a neuron across the trial by convolving the spike train with a firing rate kernel (Gaussian, = 20 ms), and then employed a function that returns a spike train whose spikes are a realization of an inhomogeneous Poisson process with the same rate profile [spike_train_generation.inhomogeneous_poisson_process(), Elephant RRID:SCR_003833, Denker 2018]. We used these rate-matched spike trains to then generate the rate-matched FNs using the methods described above.

### Dimensionality Reduction and visualization

We vectorized each FN adjacency matrix and used the UMAP algorithm to reduce the dimensionality of the FN adjacency matrix from N-by-N (neurons, N: Rs: N=143, and Rj: N= 78) to two dimensions (McInnes et al., 2018). For the full trial FNs we used the following UMAP parameters: n_neighbors=50, min_dist=1, metric=’cosine’. In Figure 2 A-D, each point in the space is an FN computed from an entire trial (total samples: Rs = 391, Rj = 246). We used a bivariate kernel density estimation for the (x,y) positions of the FNs to visualize the boundaries of the projected FNs for each reach direction. As with the full trial FN, we embedded all the temporal FNs together but in this case each sample is a FN computed from 200ms in the trial (total samples: Rs = 103246, Rj = 20856). For the temporal FNs, because there are significantly more samples, we used a different set of parameters in order to balance the local versus global structure of the data accordingly (n_neighbors=100, min_dist=0.1, metric=’cosine’). We then used a spline to interpolate a path through the set of embedded FNs from a single trial to visualize the trajectory of the temporal FN in the low-dimensional subspace (Figure 3).

### Graph Alignment Score

Similarity between two FNs, M and N with k neurons, was measured using a node-identity preserving graph alignment score, GAS, as described in Levy et al. (2020) and Gemmetto et al. (2016):

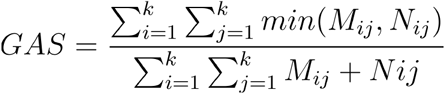

Alignment scores were grouped according to the degree difference between the instructed target in a trial. Specifically, we evaluated the GAS distributions for FNs from the same (Δ0°), neighboring (Δ45°) and opposite (Δ180°) reach directions. For the temporal networks, we computed the alignment scores between pairs of FNs within the same 200ms time window aligned to the trial cues (Instruction and Go), and to movement onset.

### Decoding

A multilayer perceptron classifier (MLPC) was trained to decode the target direction from either the FN, the firing rates, both the FNs and firing rates, or the rate-matched Poisson FNs. The MLPC architecture has one hidden layer with 100 rectified linear activation units and a constant learning rate (step size = 0.001). The MLPC was trained for 200 iterations or until convergence, whichever happened first. We trained a different decoder for each time point and feature set (75% of the data is used for training), and tested on held out data (remaining 25%). We trained and tested 50 decoders for each feature set in order to get a distribution of performance scores (Figure 5 and Supplementary Figure 1)

### Reciprocity

We computed the weighted reciprocity of FNs using a method described in Squartini et al. (2013). First, each FN was thresholded according to edge weight to isolate the most reliable interactions (thresholds indicated in the legend of Figure 7 and Supplementary Figure 2). In order to separate reciprocal weighted edges from unidirectional edges, we take the sum of the minimum weight between neuron i and j:

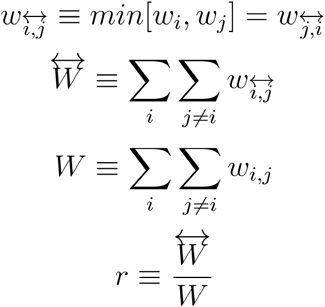

We normalized the value to the average reciprocity of 45 rate-matched networks 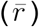 using the equation described in Garlaschelli and Loffredo (2004) for binary networks and adapted by Squartini et al. (2013) for weighted networks:

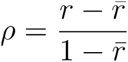

Thus when the value is positive, the FNs from the data are more reciprocal than what is expected from chance, and negative when it is less reciprocal.

## SUPPLEMENTAL INFORMATION

**Supplementary Figure 1:**
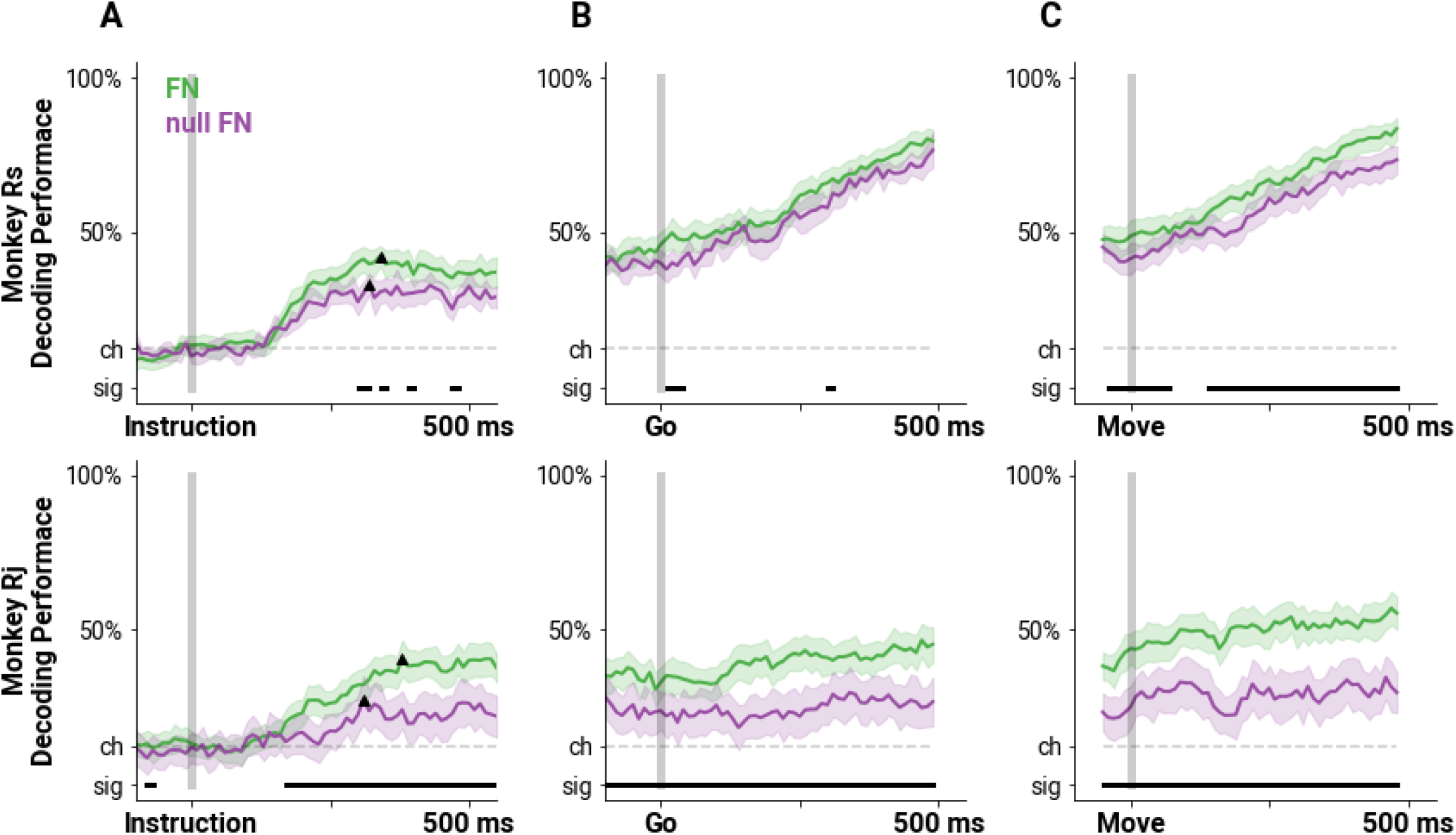
The same as Figure 5 but for decoders trained on FNs (FN, green), and rate-matched FNs (null FN, purple)

**Supplementary Figure 2:**
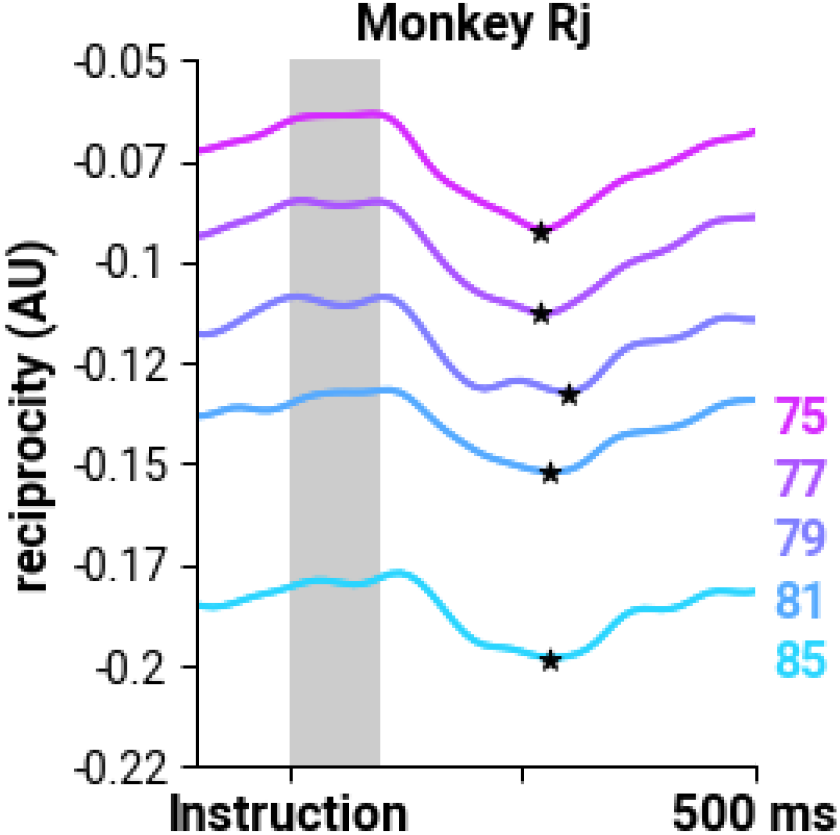
Significant decrease in reciprocity in Rj between upper quartile of edge weights. Same as Figure 6 but for edge weights from the upper quartile and above.

